# Brain-based ranking of cognitive domains to predict schizophrenia

**DOI:** 10.1101/390179

**Authors:** Teresa M. Karrer, Danielle S. Bassett, Birgit Derntl, Oliver Gruber, André Aleman, Renaud Jardri, Angela R. Laird, Peter T. Fox, Simon B. Eickhoff, Olivier Grisel, Gaël Varoquaux, Bertrand Thirion, Danilo Bzdok

## Abstract

Schizophrenia is a devastating brain disorder that disturbs sensory perception, motor action, and abstract thought. Its clinical phenotype implies dysfunction of various mental domains, which has motivated a series of theories regarding the underlying pathophysiology. Aiming at a predictive benchmark of a catalogue of cognitive functions, we developed a bottom-up machine-learning strategy and provide a proof of principle in a multi-site clinical dataset (n=324). Existing neuroscientific knowledge on diverse cognitive domains was first condensed into neuro-topographical maps. We then examined how the ensuing meta-analytic cognitive priors can distinguish patients and controls using brain morphology and intrinsic functional connectivity. Some affected cognitive domains supported well-studied directions of research on auditory evaluation and social cognition. However, rarely suspected cognitive domains also emerged as disease-relevant, including self-oriented processing of bodily sensations in gustation and pain. Such algorithmic charting of the cognitive landscape can be used to make targeted recommendations for future mental health research.

## Introduction

Schizophrenia is among the most severe mental disorders but has so far evaded mechanistic understanding. This major psychiatric disorder affects ~1% of the world population (McGrath et al., 2008)and presents a long-enduring clinical course in many patients (Hegarty et al., 1994), including social and occupational dysfunctions (Tandon et al., 2013). The associated economic costs per year range between US$94 million and US$102 billion per country (Chong et al., 2016). Schizophrenia thus imposes a huge burden on the affected individuals, their families, and society at large (Charlson et al., 2018; Wittchen et al., 2011). To eventually improve clinical care and intervention, it will be instructive to systematically explore the nature of the disease.

The clinical presentations of schizophrenia strongly suggest various cognitive impairments ranging from basic sensory perception, motor action, affective response to higher-order cognition, and social interaction (Javitt and Freedman, 2015; Taylor et al., 2012; Tost and Meyer-Lindenberg, 2012). The advent of *in-vivo* neuroimaging has enabled the investigation of the neural basis of these cognitive functions and their aberrations in disease. For more than 20 years now, functional neuroimaging experiments have accumulated hints about the candidate disease processes in schizophrenia, including for instance impaired auditory change detection (Erickson et al., 2016; Umbricht and Krljes, 2005), emotional face recognition (Kohler et al., 2010; Li et al., 2010), and working memory (Forbes et al., 2009; Schneider et al., 2007). Yet, today it is still incompletely understood “where schizophrenia is located in the brain” (Dhindsa and Goldstein, 2016; Elert, 2014; Sullivan, 2012; Weinberger and Radulescu, 2016).

Carefully designed experimental studies require that the participants attend to and execute the presented tasks for extended periods of time. The maintenance of controlled cognitive sets has sometimes been challenging to ascertain in psychiatric patients (Eickhoff and Etkin, 2016; Weinberger and Radulescu, 2016). Fortunately, mounting evidence suggests that many of the characteristic neural activity patterns described during defined experimental tasks have some correspondence in neural activity observed during task-free resting-state scanning (Bzdok et al., 2016; Cole et al., 2014; Smith et al., 2009; Tavor et al., 2016). Therefore, response-independent brain scans in clinical populations might provide unprecedented insights into brain systems dedicated to different mental operations. Additionally, despite many successes, experiments in patients with schizophrenia that test hypotheses regarding cognitive processes can carefully probe only a limited number of brain systems at a time. Such circumscribed research efforts could be complemented by computational modeling approaches that simultaneously inspect a diverse collection of cognitive functions.

The domain-transcending character of schizophrenia lends itself particularly well to take a step back and pool across a set of otherwise isolated cognitive studies. By implementing existing neuroscientific knowledge into a bottom-up pattern-learning strategy, we aimed at delineating the relevance of diverse cognitive functions in schizophrenia. An established description system of mental operations and a multi-site collection of brain scans together enabled us to derive a brain-informed ordering of the cognitive processes according to their predictive value in patients. The cognitive taxonomy has previously been used to systematically annotate roughly a quarter of the published neuroimaging literature (Derrfuss and Mar, 2009). Quantitative meta-analysis of the gathered neuroimaging resources allowed us to synthesize a large body of evidence on functional brain organization into a quintessential set of statistically defensible network-cognition associations, henceforth ‘cognitive meta-priors’. We designed and deployed a novel machine-learning approach to benchmark the brain-behavior primitives for their usefulness in distinguishing between schizophrenia patients and controls. Integrating the extracted predictive patterns of each meta-prior into a computational assay, we obtained a stratification of cognitive dysfunctions in schizophrenia. Our analytical strategy was impartial in that cognitive meta-priors of varying cortical spread were put on a comparable scale to detect their importance in the major psychiatric disorder as measured by common magnetic resonance imaging (MRI) techniques. The ranking of cognitive meta-priors was based on combined neurobiological information from brain morphology and spontaneous functional fluctuations to enable a greater generality of the results. By minimal dependence on pathophysiological, neurobiological, and statistical assumptions, we automatically computed, validated, and ranked an exhaustive set of candidate processes for their relative impairment in a major psychiatric disorder.

## Results

### Constructing cognitive meta-priors from a large-scale neuroimaging database

We jointly screened an array of cognitive functions for their predictive value in schizophrenia assisted by two complementary BrainMap taxonomies (www.brainmap.org; Fig. 1a): mental domains (Fig. 2) and experimental tasks (Supplementary Fig. 1). For each cognitive category of a taxonomy, we used coordinate-based meta-analysis to quantify the robust neural activity changes observed in thousands of healthy individuals from the BrainMap database (Fig. 1b). We generated whole-brain maps of statistically significant topographical convergence for each of 34 mental domains and 50 experimental tasks. On average, the cognitive meta-priors underlying the mental domains quantitatively synthesized 265 (ranging from 50 to 1123) database experiments, whereas the neurocognitive primitives associated with experimental tasks synthesized 167 (ranging from 50 to 701) neuroimaging studies in BrainMap (Supplementary Tables 2 and 3). As expected, the cognitive concepts clearly differed in their distributed set of responsive brain regions.

**Figure 1.**
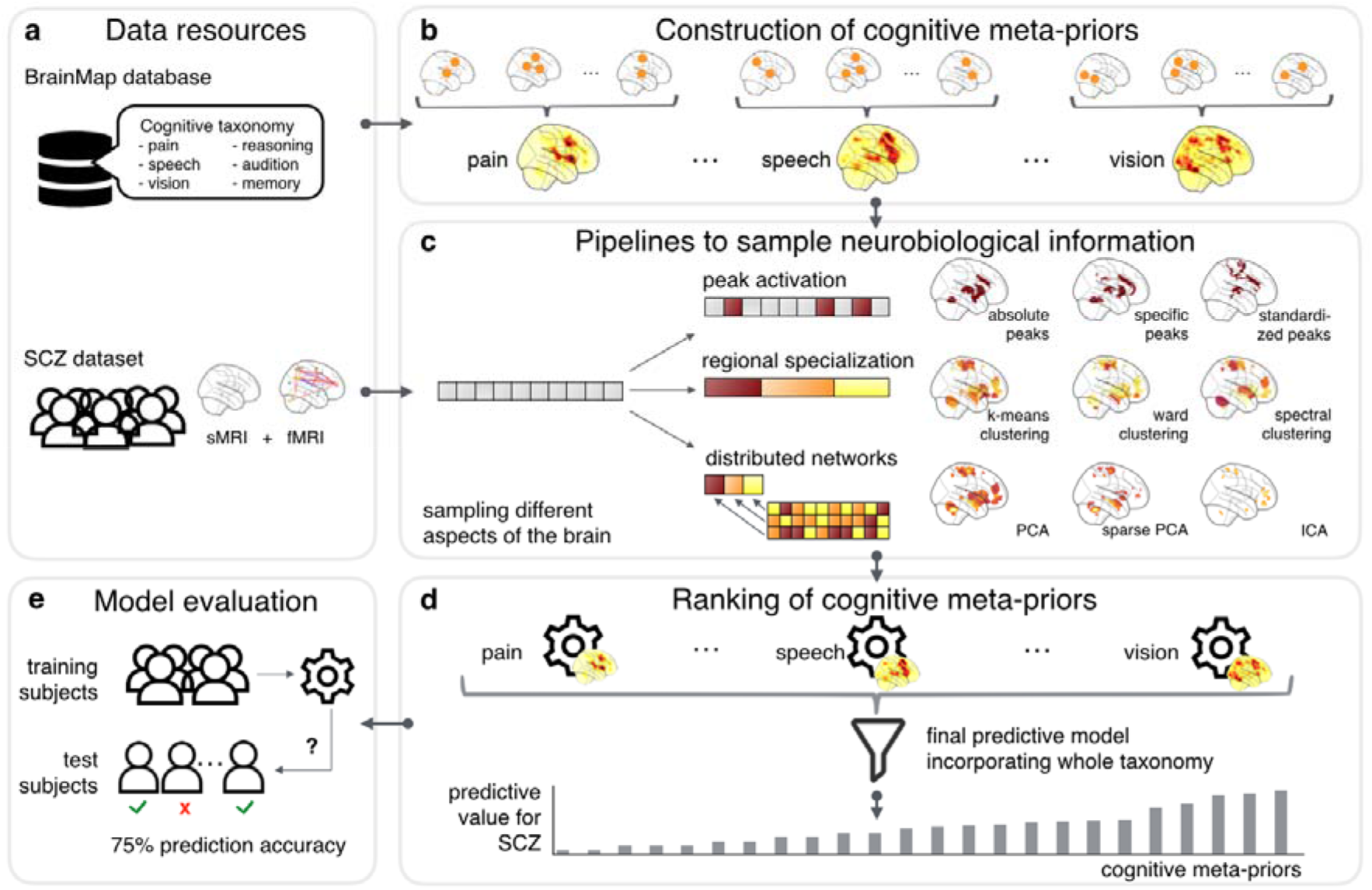
Analysis workflow. Illustrates our tactic for bottom-up ranking candidate cognitive processes for playing a relevant role in schizophrenia (SCZ) disease, **(a)** The large neuroimaging database hosts consistent neural activity findings annotated with a detailed taxonomy of cognitive domains. The pre-existing neuroscientific knowledge was capitalized on for analyzing a multi-site dataset of patients with schizophrenia and healthy controls. **(b)** Meta-analytic networks summarized each cognitive process (e.g., pain) from the neuroimaging database. **(c)** The cognitive meta-priors guided nine pipelines to extract complementary neurobiological characteristics from structural and functional magnetic resonance imaging data (sMRI and fMRI) of the schizophrenia dataset. **(d)** Each cognition-network pair was put to the test of telling patients and controls apart. Brain-driven predictions of all meta-priors were combined in a higher-level summary model that compared cognitive processes in their usefulness for disease classification. Ensuing relevance scorings of cognitive processes were averaged across nine complementary sampling pipelines. The cognitive-domain-spanning final classifier thus charted the landscape of altered cognitive processes in schizophrenia, while ensuring each meta-prior to have the same opportunity. **(e)** Success in disambiguating patients vs. controls was confirmed by explicit validation of the predictive model in brain maps from previously left-out participants (10-fold cross-validation scheme).

Regarding mental domains (Fig. 2; see Supplementary Fig. 1 for experimental tasks), a database of studies linked to motor inhibition, for example, consistently engaged mainly the bilateral supplementary motor area and posterior insula, but also involved the putamen, frontal eye field, and intraparietal sulcus in most individuals. Working memory processes, instead, robustly recruited a distributed set of bilateral brain regions, including the dorsolateral frontal cortex and dorsal anterior cingulate cortex but also the inferior parietal lobe and precuneus. The neural responses coherently observed across experiences of fear were located in the bilateral amygdala, extending into the neighboring hippocampus as well as bilateral anterior cingulate cortex, and ventromedial frontal cortex and the right posterior insula. The neural activity pertaining to visual perception was prominent in the bilateral inferior and middle occipital gyri of the early visual cortex, extending into the fusiform gyri, and posterior parietal lobe, as well as middle and inferior temporal gyri (not shown). For both BrainMap taxonomies, the cognitive meta-priors encapsulated quantitatively dissociable patterns of whole-brain activity underlying a class of cognitive processes.

**Figure 2.**
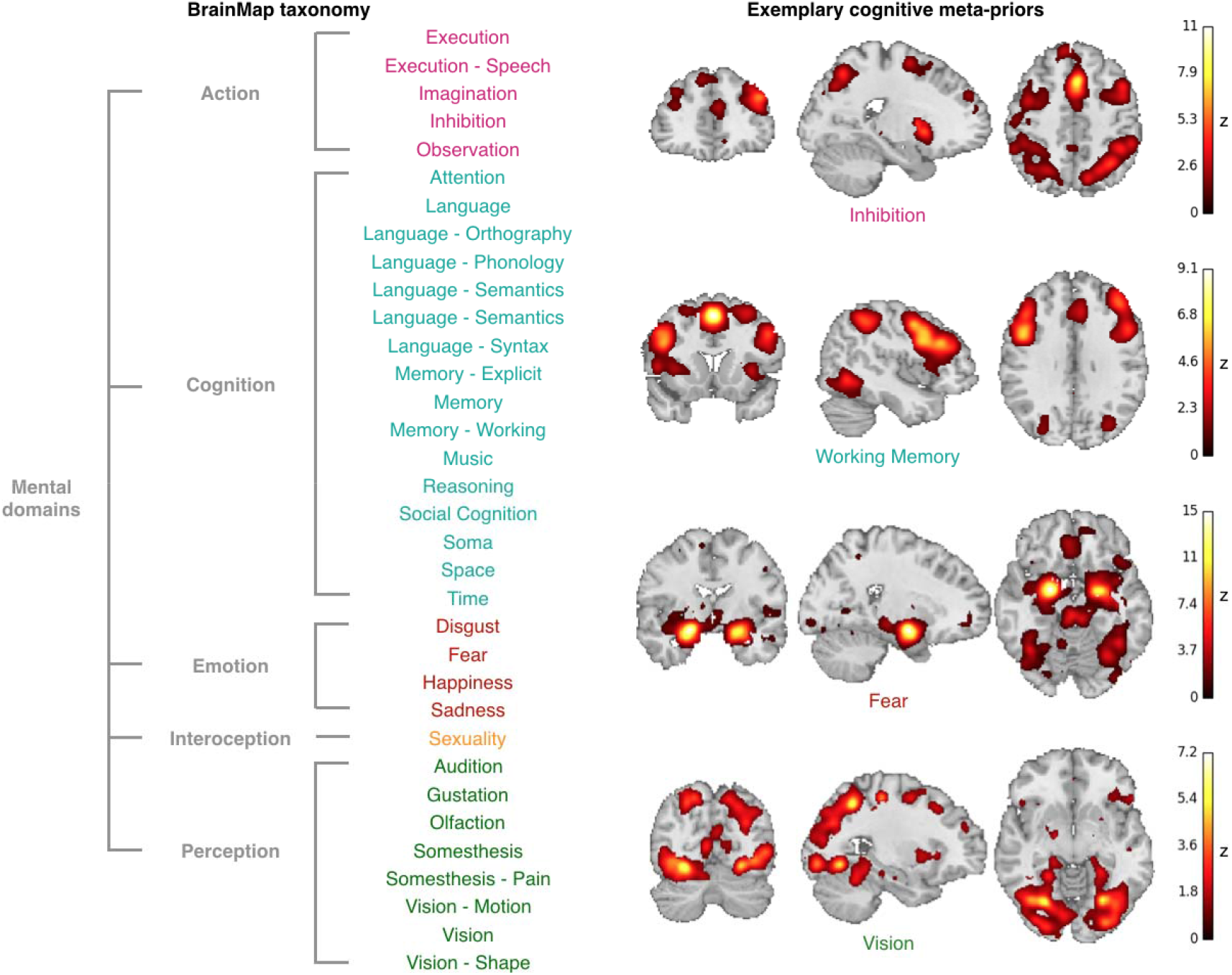
Overview of a taxonomy that compartmentalizes human cognition. (Left) Exhaustive set of mental operations used for brain-driven ranking of altered cognitive concepts in schizophrenia. BrainMap defines two description systems: mental domains (shown here) and experimental tasks (Supplementary Fig. 1). Note that the five top classes (Action, etc.) were disregarded in the present study to avoid hierarchical dependence between the cognitive classes. Database offers results of almost a quarter of the published functional neuroimaging experiments carefully annotated with both taxonomies. **(Right)** A cognition-topography map for each cognitive category was generated from the neuroimaging database. Four examples of cognitive meta-priors are shown (z-scored for display, only voxels with positive z-scores shown).

We made several general observations across the synthesis of neurocognitive priors underpinning mental domains and experimental tasks. Various cognitive processes related to basic perception corresponded to bilateral activity increases in primary sensory and association cortices and insula, but also mapped onto the putamen as well as superior and medial frontal gyri. Different cognitive processes associated with motor action primarily elicited bilateral activity increases in the precentral gyri and thalamus, and also recruited insula and medial frontal gyri. Emotion-related processes were mainly subserved by the bilateral amygdala and both anterior and posterior cingulate cortex but also frequently involved the medial prefrontal cortex. Many higher-level cognitive processes tended to predominantly evoke increases in neural activity in bilateral prefrontal regions and posterior cingulate cortex, and further involved bilateral inferior and superior parietal lobe and middle temporal gyri. Across domains of both taxonomies, mental domains and experimental tasks, we observed that many cognitive functions were underpinned by a distributed constellation of higher- and lower-level brain systems.

### Estimating model performance across imaging modalities and brain sampling approaches

The neurocognitive primitives that we obtained from each taxonomy guided the extraction of structural (sMRI) and functional (fMRI) brain measurements in a five-site schizophrenia dataset (n=324, mean age=35.4 ± 11.5 years; Table 1). We retrieved the neurobiological information using nine complementary brain sampling approaches revolving around neural activity changes i) at peak locations, ii) in brain regions, and iii) distributed brain networks in a given cognitive class (Fig. 1c). These reshaped brain data were used to build one predictive pattern-learning model for each cognitive meta-prior. That is, one trained linear classifier tracked one particular element from a taxonomy when applied to the whole-brain sMRI and fMRI data to separate patients from controls. These domain-specific classifiers only learned from brain information that was shown to be linked to a particular mental operation across hundreds of neuroimaging experiments. The domain-specific predictive models served as components for a taxonomy-overarching predictive model (Fig. 1d). Initially, we wished to assess the ability of the summary model to prospectively distinguish between patients with schizophrenia and healthy controls by repeatedly testing the predictive model in held-out participants who were not seen by a pattern-learning algorithm (10-fold cross-validation; Fig. 1e). Across brain structure and function and complementary brain sampling tactics, the domain-integrating summary model performed consistently better than chance (50%) in classifying new individuals (Supplementary Fig. 2).

**Table 1:** Clinical sample from different sites. Patients with schizophrenia (SCZ) and matched healthy controls (HC); age values in mean ± standard deviation; ^1^ statistical comparison of age differences between groups performed via t-test; ^2^ statistical comparison of sex differences between groups performed via chi-squared test.

| **Sample** | **n**<br>**(males)** | **Sex differences**<br>**(p-values) <sup>1</sup>** | **Age**<br>**(years)** | **Age differences**<br>**(p-values) <sup>2</sup>** |
| --- | --- | --- | --- | --- |
| *Groningen* |  |  |  |  |
| SCZ | 32 (19) |  | 33.6 ± 11.1 |  |
| HC | 32 (19) | 0.476 | 31.6 ± 11.2 | 1 |
| *Göttingen* |  |  |  |  |
| SCZ | 32 (26) |  | 32.3 ± 9.9 |  |
| HC | 29 (22) | 0.889 | 31.9 ± 9.4 | 0.841 |
| *Aachen* |  |  |  |  |
| SCZ | 14 (11) |  | 35.1 ± 11.1 |  |
| HC | 13 (10) | 0.682 | 33.2 ± 12.0 | 0.719 |
| *Lille* |  |  |  |  |
| SCZ | 15 (9) |  | 33.3 ± 5.0 |  |
| HC | 16 (11) | 0.048 | 29.0 ± 6.3 | 0.894 |
| *COBRE* |  |  |  |  |
| SCZ | 68 (55) |  | 38.2 ± 13.7 |  |
| HC | 73 (50) | 0.270 | 35.8 ± 11.7 | 0.136 |
| ***Total analyzed sample*** |  |  |  |  |
| SCZ | 161 (120) |  | 35.4 ± 11.9 |  |
| HC | 163 (112) | 0.125 | 33.4 ± 11.0 | 0.299 |

We first compared the classification performance across imaging modalities (Supplementary Fig. 2). We observed that predictive models only informed by brain structure correctly predicted disease state in 73.2% (SD=2.1%, across brain sampling approaches) of new individuals for mental domains and in 73.5% (SD=1.9%) for experimental tasks on average. Predictive models aware of interindividual differences in brain structure outperformed predictive models based on functional and combined imaging modalities in 4 of 9 brain sampling approaches in mental domains and in 5 of 9 approaches in experimental tasks. We further observed that the average classification accuracy of predictive models of meta-priors informed by brain function reached 70.9% (SD=1.6%) for mental domains and 70.8% (SD=1.4%) for experimental tasks. The predictive models that only had access to brain function were found to be superior to structural and combined imaging modalities in 1 of 9 brain sampling approaches in mental domains and in 0 of 9 brain sampling approaches in experimental tasks. In combined brain structure and function, we found mean classification performances of 73.4% (SD=1.9%) in mental domains and 73.6% (SD=2.1%) in experimental tasks. The predictive model tuned to combined information from brain structure and brain function performed better than predictive models based on a single imaging modality in some but not all brain sampling approaches in both mental domains and experimental tasks. Across different types of brain information, these slight differences in classification performance were not statistically significant at p<0.05 as indicated by our bootstrapped 95% confidence intervals. On average, however, predictive models utilizing both brain structure and brain function achieved the highest classification performances.

We then examined the outcome of different brain sampling approaches (Supplementary Fig. 2). We observed varying prospective classification performances depending on the type of brain data. In combined brain structure and function, for instance, the classification performance ranged from 69.8% ([57.6%; 81.3%], bootstrapped 95% CI) to 75.9% ([62.5%; 84.4%]) across mental domains. Among experimental tasks, in turn, the prediction accuracy ranged from 69.8% ([60.6%; 81.3%]) to 75.9% ([68.8%; 87.9%] across different data-extraction approaches. Comparing the different brain sampling approaches, different activation-, region- or network-focused strategies were advantageous in different settings for separating patients from controls, without a consistent winner.

It is important to note that our analyses were based on brain data after adjusting for age, sex, and data acquisition site to prevent the predictive model from capturing variation due to variables of no interest. In brain data without this confound removal step, we observed instances of slightly elevated classification performance of the predictive model. This piece of evidence suggests that the predictive model learned useful information from nuisance variables. Nevertheless, the uncorrected sub-analyses yielded a comparable ranking order, which provided evidence that the relevant brain variation was mostly independent of age, sex, and site.

### Testing the cognitive specificity of schizophrenia predictability

To further ensure that the pattern-learning algorithms captured the individual compositions of cognitive facets instead of potential confounding influences, we validated the top-level predictive model in a negative test. We formally compared the classification performance of the domain-spanning model to a cognition-naïve null model. In 1000 random permutations, we specifically permuted how domain combinations co-occurred within individuals of the health or disease group. By leaving all other data characteristics intact, we wished to reject the null hypothesis that the constellation of cognitive functions of the participants were not relevant for group classification. The non-parametric hypothesis test revealed that the summary model discriminated between patients and controls significantly better than the null model across brain sampling approaches at a level of p<0.05 for mental domains and p<0.01 for experimental tasks (Fig. 3). That is, we observed our actual or a higher prediction accuracy in less than 50 of 1000 cases in mental domains and in less than 10 in 1000 cases in experimental tasks if there was no systematic relation between an individual’s cognitive relevance and group detection. In short, our negative test ascertained that the successful classification performance of the composite predictive model could be defensibly ascribed to the participant-specific configurations of cognitive aspects.

**Figure 3.**
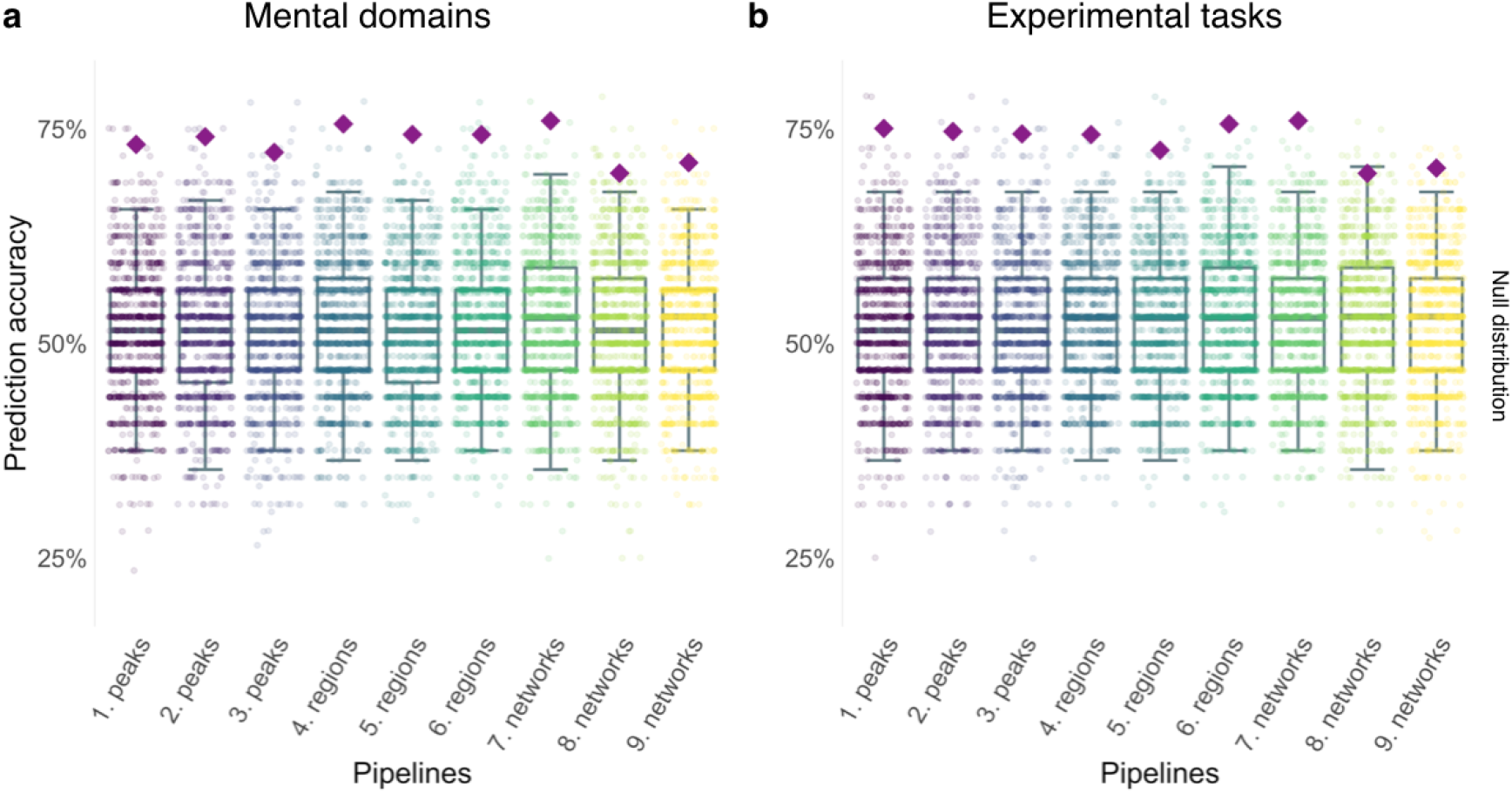
Validation of our data-analysis framework. The final predictive model classified healthy versus schizophrenic individuals statistically significantly better than a cognition-naive null model in each of two taxonomies. We estimated the null distribution by selectively corrupting the participant pattern of cognitive indices while leaving other structure in the data intact. To ascertain that the final predictive model captured participant-specific cognitive facets instead of confounding variables. Purple diamonds indicate the (out-of-sample) classification performance of the composite model based on mental domains **(a)** and experimental tasks **(b)** using combined structural (sMRI) and functional (fMRI) brain information. The dots show 1000 model performances realized under the null hypothesis. The grey boxplots show bold lines for median (50^th^ percentile), the lower and upper quartile (25^th^ and 75^th^ percentile), and whiskers for the interquartile distance (25^th^ - 75^th^ percentile) besides the box. In each of nine ways to sample brain information, the composite model performed significantly better than the null model (p < 0.05 for mental domains and p < 0.01 for experimental tasks). If the individual combinations of cognitive expressions were not relevant, we would only observe our actually obtained prediction performance (purple diamond) in at most 50 out of 1000 cases for mental domains and in at most 10 out of 1000 cases for experimental tasks. The negative test implies that the successful individualized decisions of our predictive model can be ascribed to participant-specific cognitive alterations rather than other characteristics of the participant sample.

### Determining contributions to schizophrenia predictability across cognitive domains

The domain-spanning summary model ensured the comparability of the cognitive classes despite naturally varying numbers of distributed brain regions that constituted a cognitive brain network. That is, the top-level predictive model enabled us to impartially contrast the cognitive meta-priors in their relative contribution to schizophrenia classification in each type of brain data.

Informed by both brain structure and function, the rankings of mental domains (Fig. 4) and experimental tasks (Fig. 5) clearly demonstrated that some candidate processes were often more relevant for schizophrenia detection than others. Regarding mental domains, for instance, the top-scoring domains of gustatory perception, pain perception, and experience of sadness were found to be statistically significantly more predictive of schizophrenia than attention, temporal reasoning, and visual perception at p<0.05 (Fig. 4). Regarding experimental tasks, mental processes related to pain discrimination, face discrimination, and visual tracking were significantly more discriminable of disease state than the lowest ranked mental operations underlying listening to and producing music, visuospatial attention, and the Stroop task at a level of p<0.05 (Fig. 5). Across both taxonomies, we observed that cognitive functions related specifically to social-affective (e.g., experience of sadness and face discrimination tasks) and internally oriented perception processes (e.g., pain perception and pain discrimination tasks) emerged as most critical for schizophrenia classification. Thus, we quantitatively identified common and distinct elements of cognition in MRI-based imaging of brain structure and function in their utility for the study of schizophrenia.

**Figure 4.**
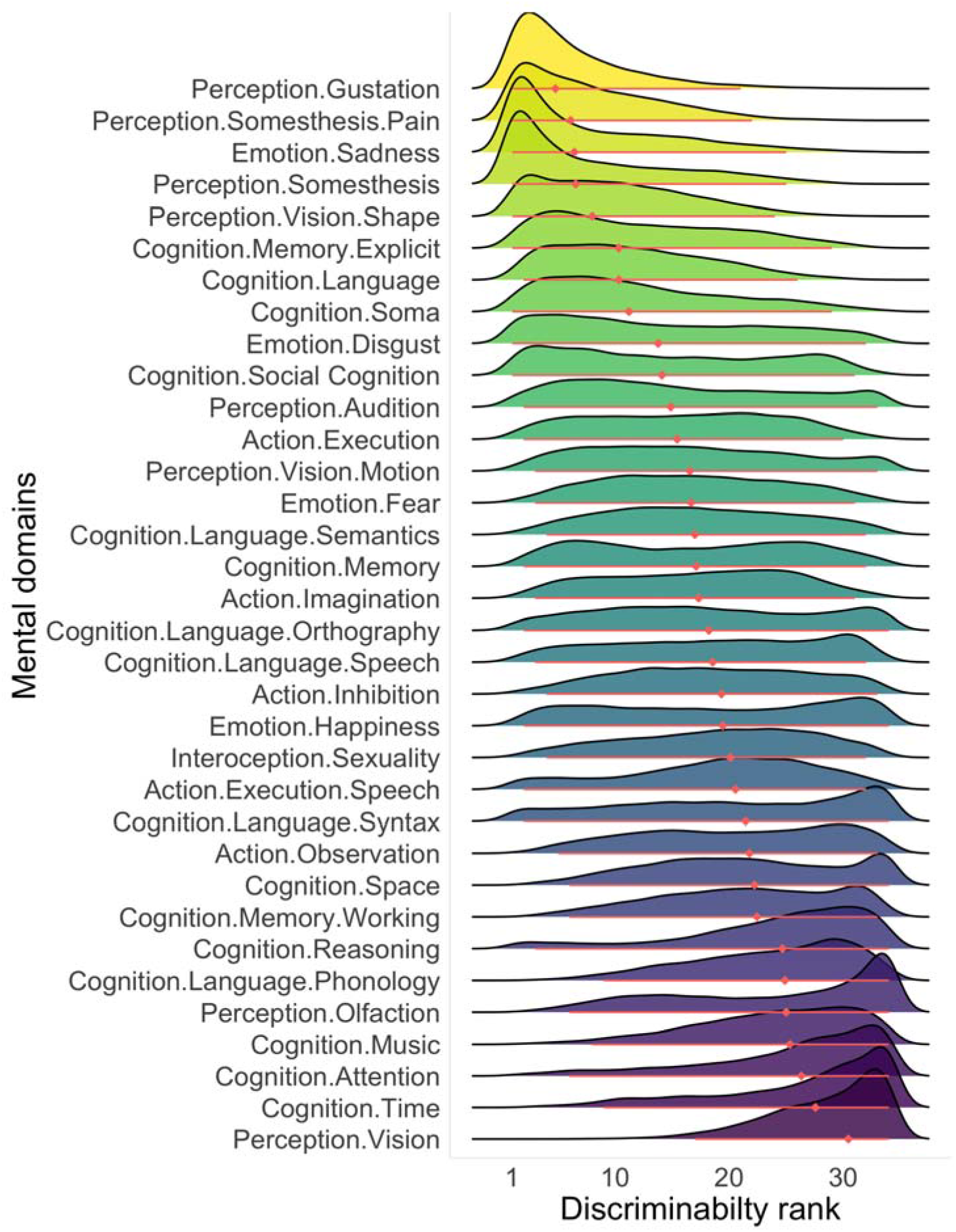
Quantified predictive value of mental domains in schizophrenia. We systematically screened for dysregulated cognitive processes to facilitate the development of personalized diagnoses and new treatment strategies. Relative contribution of mental domains in disambiguating patients with schizophrenia and healthy controls. 34 mental domains ordered according to their average ability to forecast disease state. Joyplot shows weighted importance ranks for each domain (colored mountains). Red diamonds depict mean ranking position across brain sampling strategies (see Supplementary Fig. 3 for pipeline-specific domain ranks). Certainty of discriminability position was assessed by estimating bootstrapped 95% population intervals (red lines). For instance, gustation was highly predictive across complementary approaches to sample neurobiological information, whereas the relevance of audition was more depended on the sampling pipeline. Some intensively studied concepts of attention (e.g., Braff, 1993) and working memory (e.g., Forbes et al., 2009; Lee and Park, 2005) have been situated among the cognitive classes least predictive for schizophrenia. All results based on combined sMRI and fMRI data.

**Figure 5.**
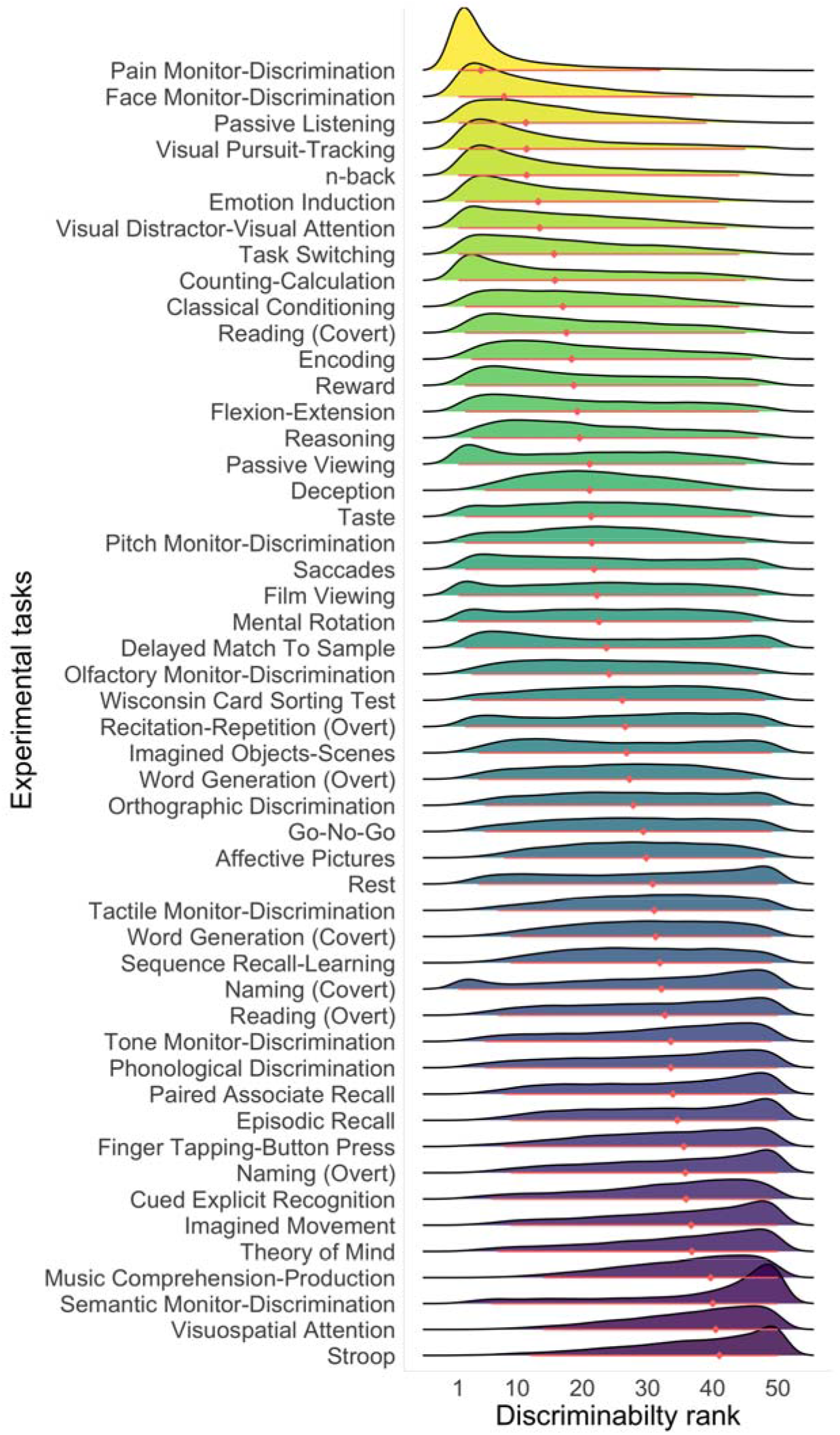
Quantified predictive value of experimental tasks in schizophrenia. We broadly screened for distinctive experimental paradigms to facilitate the development of personalized diagnoses and new treatment strategies. Relative contribution of experimental tasks in disambiguating patients with schizophrenia and healthy controls. 50 experimental tasks were ordered according to their average ability to forecast disease state across brain sampling strategies (see Supplementary Fig. 4 for pipeline-specific domain ranks). Joyplot shows weighted importance ranks for each domain (colored mountains). Red diamonds depict mean position in relevance. Certainty of discriminability position was assessed by estimating 95% population intervals (red lines). Precision estimates computed by repeatedly resampling participants with replacement (bootstrapping). All results based on combined sMRI and fMRI data.

We further elaborated this evidence from combined brain structure and function in predictive models that drew on a single imaging modality. In brain volume alone, 7 of the top 10 mental domains were in agreement with the top 10 mental domains found in combined sMRI and fMRI data. However, knowledge of speech, sensing sexual needs, and speaking were ranked among the top 10 mental domains in brain structure instead of pain perception, knowledge of language, and body knowledge. Similarly, 5 of the top 10 ranked experimental tasks in brain structure were also counted among the top 10 experimental tasks in combined brain structure and function. Yet, mental processes elicited by film viewing, encoding, theory of mind, passive viewing, and Wisconsin Card Sorting Test tasks emerged as more disease-relevant in brain structure than mental operations involved in visual tracking, visual attention, task switching, numerical operations, and classical conditioning experiments. In intrinsic functional connectivity alone, 6 of the top 10 mental domains corresponded to the top 10 mental domains in combined sMRI and fMRI data. However, motor execution, motion perception, experience of fear, and auditory perception were counted among the top 10 mental domains in brain function instead of knowledge of language, body knowledge, experience of disgust, and social cognition. Similarly, 7 of the top 10 ranked experimental tasks in brain function were concordant with the top 10 found in combined brain structure and function. Here, mental operations related to flexing and extending movements, rapid eye movements, and passive viewing tasks arose as more discriminable of schizophrenia in brain function than mental processes elicited by face discrimination, task switching, and numerical operations paradigms. Despite several modality-specific relevances of cognitive domains in schizophrenia, the obtained utility rankings were largely overlapping based on different types of brain data.

### Isolating non-linear predictive relationships across cognitive domains and predictive relevance maps

Finally, we wished to explore particular ways in which domain pairs act together in distinguishing between patients and controls (Fig. 6). For this purpose, we estimated the non-linear interaction surface of cognitive domain pairs in classification of disease state after accounting for the effects of the remaining domains of the taxonomy (i.e., partial dependence estimation). The results demonstrated that beyond-linear relationships contributed to how specific pairs of cognitive processes allowed us to distinguish between patients and healthy controls. Charting these two-way interactions of the most predictive mental domains and experimental tasks showed various types of links between schizophrenia classification and single cognitive concepts. Besides approximately linear links to schizophrenia prediction (n-back task and passive listening), we also observed somewhat logarithmic (experience of sadness), exponential (pain perception and pain discrimination), and polynomial (gustatory perception and face discrimination) non-linear relationships. Similarly, we found different qualities of relationships such as of approximately linear (semantic discrimination and Stroop task), logarithmic (visual perception), exponential (temporal reasoning), and polynomial kind (listening to and producing music) among the lowest ranked domains (Supplementary Fig. 5). The comparison of the more-than-linear effects of two top-ranked and two less successful domains also showed that schizophrenia classification relied more heavily on statistical dependencies among the top ranked domains as compared to the lowest ranked domains. When we directly contrasted top and lowest ranked domains, we again found stronger contributions of the top ranked domains on the classification of health versus disease compared to the lowest ranked domains (e.g., gustatory perception vs. attention, and face discrimination vs. Stroop task). Across imaging modalities and cognitive description catalogues, we observed that cognitive functions contributed in complex ways to schizophrenia classification, that is, patterns in brain data to which purely linear Pearson correlation and regression-type analyses are blind.

**Figure 6.**
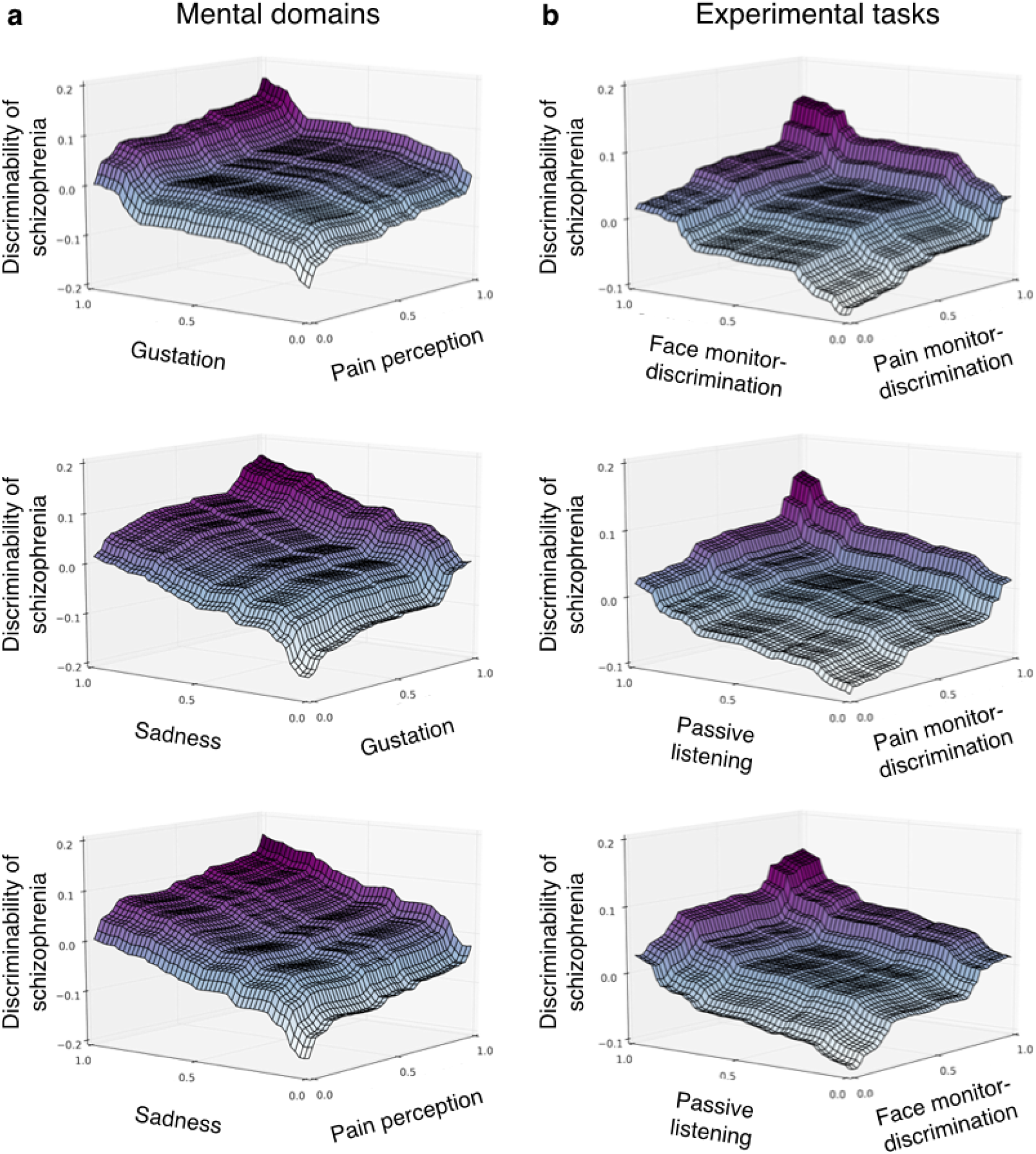
Domain-domain interactions in detecting schizophrenia. Inspects in more detail potentially complicated relationships in the predictability of mental domains **(a)** and experimental tasks **(b)** after accounting for influence of the remaining domains (instead of ignoring them). Partial dependence of schizophrenia predictability (z-axis) on the joint distribution of two selected cognitive domains (x- and y-axis) in predicting whose brain scans are from a schizophrenia patient (PCA pipeline). Pairs of the top 3 cognitive processes are shown for both taxonomies (see Supplementary Fig. 5 for further examples). All results based on combined brain imaging types (sMRI and fMRI data).

After contrasting the individual meta-priors for their differences, we wished to further explore common characteristics across the cognitive classes of a taxonomy in schizophrenia prediction. To examine which brain areas were most discriminative of disease state across cognitive domains, we globally mapped the importances of each cognitive meta-prior onto the brain. In both mental domains and experimental tasks, we observed largely overlapping patterns of brain regions that were most relevant for disease classification (Fig. 7). This similarity across two distinct ways to catalogue cognitive functions serves as post-hoc validation of our approach.

**Figure 7.**
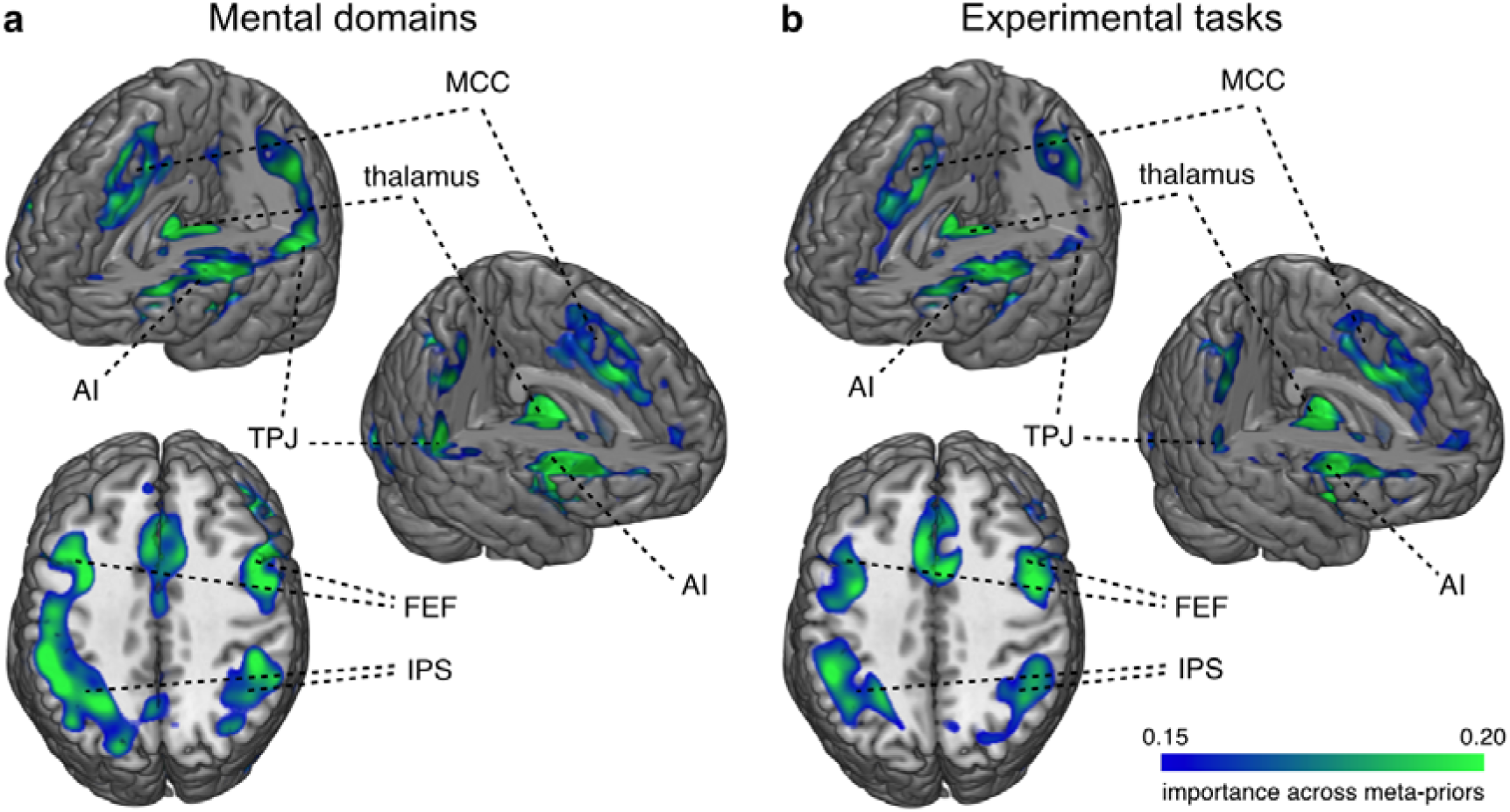
Predictive relevance maps for schizophrenia. Quantifies the average extent to which individual brain regions contributed to disease classification across 34 mental domains **(a)** and 50 experimental tasks **(b)** experimental tasks. Whole-brain maps depict relative importance values of the cognitive meta-priors across nine brain sampling strategies. In both taxonomies, nodes of the dorsal attention network (e.g., frontal eye field (FEF) and intraparietal sulcus (IPS)) and saliency network (e.g., anterior insula (AI) but not mid cingulate cortex (MCC)) as well as thalamus were highly pertinent in distinguishing patients from controls. Left and right temporo-parietal junction (TPJ), however, emerged as discriminative in mental domains but not in experimental tasks. Both taxonomies provided largely overlapping but still distinct brain patterns underlying schizophrenia classification. This converging evidence across two independent cognitive taxonomies further strengthens the validity of our approach. Brain maps were smoothed (FWHM = 6mmm) and thresholded for display (see https://neurovault.org/collections/4074/ for unthresholded predictive brain maps). All results are based on combined sMRI and fMRI data.

## Discussion

How can we derive principled recommendations for psychology and neuroscience experiments from brain recordings that can be measured at scale? We have introduced a machine-learning strategy to stratify a catalogue of cognitive classes according to their utility in identifying schizophrenia. We first distilled existing neurobiological knowledge on constituent elements of the human mind into couples of cognitive concept and quintessential neural representation. In a bottom-up fashion these cognitive meta-priors were contrasted in their capacity to distinguish between patients and controls based on easily acquired and commonly available structural and functional brain scans of a multi-site study of a large schizophrenia cohort. Each domain-specific classifier was exclusively based on brain information with a pre-established link to a specific cognitive domain. The analytical framework was impartial in giving each cognitive category the same opportunity to be selected as most important in identifying schizophrenia. The data-guided ranking highlighted certain cognitive categories that were more discriminative for this major psychiatric disorder than other types of mental activity. It is key outcome that we found both frequently investigated and largely untapped disease concepts to be relevant in schizophrenia.

Among the traditionally examined concepts, our across-systems analysis underscored the critical role of tones and speech appraisal (passive listening) as 3^rd^ most relevant among 50 experimental tasks and the sense of hearing (audition) as 11^th^ among 34 mental domains, on the one hand. This quantitative evidence from structural and functional brain scans confirms the long-standing clinical emphasis on auditory hallucinations as a hallmark symptom of schizophrenia patients. Hearing malevolent voices and other auditory misperceptions (Lim et al., 2016; Llorca et al., 2016; McCarthy-Jones et al., 2017) might be mediated by impaired pre-attentive filtering mechanisms (Javitt, 2009; Javitt and Freedman, 2015; Javitt and Sweet, 2015; Rissling and Light, 2010). Additionally, neuroimaging studies have consistently shown abnormalities of auditory brain regions in schizophrenia using meta-analyses: (i) structural neuroimaging studies found reduced volume in parts of the auditory cortex in the superior temporal gyrus (Honea et al., 2005; Modinos et al., 2013) and (ii) functional investigations have reported increased neural activity in areas related to speech perception, language, and memory during active sensations of speech hallucinations (Allen et al., 2012; Jardri et al., 2011).

A relation of auditory processing systems with language and memory was suggested by theories that ascribe the formation of auditory hallucinations to intrusive memories and the external misattribution of inner speech (Curcic-Blake et al., 2017; Jardri et al., 2011). In line with these notions, our stratified prediction experiment revealed the relevance of memory and language systems (explicit memory and language). Besides the auditory system, also visual processing (e.g., shape vision and visual tracking) emerged as disease-relevant - a sense commonly affected by hallucinations and psychosis (Waters et al., 2014). The disturbance of these basic sensory processes might lead to deficits in higher-level cognition and thus impact functional outcome in patients, which is supported by a recent structural equation modeling study (Javitt and Freedman, 2015; Thomas et al., 2017). Difficulties in the interpretation of speech prosody were argued to potentially entail problems in social interaction (Javitt and Freedman, 2015; Javitt and Sweet, 2015). Our large-scale analyses provide evidence for the potential of MRI-facilitated clinical control especially for distressing and at times dangerous voice hallucinations.

Furthermore, viewing or evaluating information from others’ faces (face discrimination) as a core element of social cognition was attributed 2^nd^ highest importance among 50 experimental tasks pooling across brain structure and function. Concurrently, emotional processing, as another important aspect of social behavior (Green et al., 2015; Ochsner, 2008), emerged as relevant for schizophrenia: appraisal of environmental cues with affective valence (emotion induction) was ranked 6^th^ among 50 experimental tasks, while the experience of two negative basic emotions, sadness and disgust, were ranked 3^rd^ and 9^th^ among 34 mental domains. The more general concept of information processing related to fellow humans (social cognition), was 10^th^ among 34 mental domains. By carrying out a quantitative cognitive screening, we endorse the broader relevance of social-affective thought and behavior in schizophrenia. This comparably recent research trend in psychiatry and neuroscience is currently gaining momentum for several reasons: (i) previous studies have hinted at a broad spectrum of social dysfunctions in patients with schizophrenia (see Savla et al. (2013) for a meta-analysis), (ii) such dysfunctions seem to be time-enduring and already present in prodromal phases of the disease (Bora et al., 2009; Green et al., 2012), and (iii) the deficits in social cognition are intimately related to poor work and community functioning (Couture et al., 2006; Fett et al., 2011; McCleery et al., 2016). Moreover, aberrations in socio-emotional processing circuits seem to mediate the impact of social environmental risk factors such as urbanization and migration on schizophrenia (Tost and Meyer-Lindenberg, 2012).

Our quantitative outcomes are also supported by earlier documentation of emotional processing disturbances in schizophrenia (Aleman and Kahn, 2005; Derntl et al., 2009; Derntl et al., 2012). Several emotion recognition studies reported larger impairments in the processing of faces with negative, rather than positive, emotions such as sadness and disgust (Kohler et al., 2010). The predominance of impaired negative emotion recognition was also reflected in our results by higher disease-discriminability of negative emotions such as sadness, disgust, and fear as opposed to the positive emotion happiness. In line with our evidence, a meta-analysis of structural MRI studies in schizophrenia highlighted gray matter decreases in amygdala and insula (Ellison-Wright et al., 2008). Another MRI study found a positive correlation between amygdala volume and performance in recognizing sad faces (Namiki et al., 2007). Furthermore, a meta-analysis of functional MRI studies revealed reduced brain activation in amygdala, parahippocampal gyrus, and fusiform gyrus, but increased activity in left insula during emotional face processing (Li et al., 2010).

The aberration of social-affective brain systems in schizophrenia also translates into various clinical symptoms. Patients often exhibit negative symptoms, such as diminished emotional expression and apathy, which tend to have enduring trajectories compared to the more episodic positive symptoms. Indeed, impairments in social cognition have been proposed to have a stronger impact on functional outcome than other cognitive impairments (Carrión et al., 2016; Fett et al., 2011). The collective outcomes of our cognitive charting underscore the potential of clinical interventions that target affective impairments in schizophrenia such as social rehabilitation and training regimens in various social skills (Kurtz and Richardson, 2012).

Besides reinforcing currently studied forms of thinking aberration, certain cognitive domains emerged as critical to schizophrenia patients that have so far seldom been the center of investigation. For instance, the appraisal of pain-related cues was ranked as the 1^st^ most predictive experimental task (pain discrimination) and 2^nd^ most affected mental domain (pain perception). Despite a documented decrease in the sensitivity of patients to pain stimuli in the clinical setting, this phenomenon has been the object of very few experimental studies (Dworkin, 1994; Stubbs et al., 2015). A recent review on the limited available literature suggested the existence of impairments in the sensory-discriminative, affective, and cognitive components of pain processing (Stubbs et al., 2015). Similarly, one of the few existing neuroimaging studies on the topic reported decreased recruitment of pain-responsive brain regions such as the anterior insula and increased recruitment of sensory-processing-tuned brain systems such as the primary somatosensory cortex during pain processing in patients with schizophrenia (de la Fuente-Sandoval et al., 2010).

Combining these streams of evidence, we used our cognitive ranking outcomes to identify a critical role of pain and emotion appraisal in schizophrenia. In combination with the highly scored domains related to processing external and internal bodily feedback such as skin sensations (somatic cognition and somesthesis), many of the most predictive cognitive classes can be considered as impaired interoceptive integration. This contention of a key role of binding bodily information is further supported by the 1^st^ rank of the sense of tasting (gustation). Indeed, misperceptions of imagined tactile or bodily cues (e.g., feeling of insects on the skin) is an often-encountered clinical symptom in schizophrenia patients (McCarthy-Jones et al., 2017; Thomas et al., 2007). More broadly, this major psychiatric disease is often considered to be a disorder of the self and subjective experience (Fletcher and Frith, 2009). Colloquially, these patients may suffer from a misbalance in “how the body listens to itself” - how it senses, integrates, and prepares reactions to somatic signals. Alterations of interoceptive capacity might also give rise to misattribution of inner signals to external sources, which is a frequently observed clinical feature of patients with schizophrenia. Dysregulated processing of nociceptive and autonomic signals has very recently been raised as a potential mechanism involved in schizophrenia pathophysiology and potentially other mental disorders (Ardizzi et al., 2016; Khalsa et al., 2017; Owens et al., 2018). More generally, pain insensitivity can lead to poor help-seeking behavior. Hence, future research endeavors enlightening pain processing in schizophrenia might help to reduce the high morbidity and mortality observed among schizophrenia patients (Dworkin, 1994; Stubbs et al., 2014; Stubbs et al., 2015). Both pain appraisal and gustation are exemplary instances of the used taxonomy that share the processing of internal bodily information and associated feedback loops. Taken together, the identification of these highly discriminative cognitive classes stresses the research potential of elucidating how bodily signals from the internal organs which may expose a currently underappreciated disease mechanism.

This interpretation is in line with another quantitative finding of a yet mostly unexplored domain. Tasting or imagining the flavor of food reached the 1^st^ position among 34 candidate mental domains which probably relates to epidemiological studies that estimate 7% to 31% of patients with schizophrenia experience some form of gustatory hallucination (Baethge et al., 2005; Connolly and Gittleson, 1971; Lewandowski et al., 2009; Thomas et al., 2007). The scarce studies directly examining gustation in patients with schizophrenia reported a significant deficit in their sensitivity for different tastes (Balderston et al., 2003), such as the bitter-tasting antiheroic compound phenylthiocarbamide (Moberg et al., 2007; Moberg et al., 2005). Additionally, there is some tentative evidence for abnormalities in brain regions related to gustation including the insula, thalamus, and orbitofrontal cortex (Balderston et al., 2003). For instance, taste chemoreceptors responses seem to be reduced in the dorsolateral prefrontal cortex (Ansoleaga et al., 2015). The subordinate role of gustatory hallucinations in common clinical assessments, as opposed to other sensory misperceptions, may be one reason why research on disturbed gustatory processing in schizophrenia may still be in its infancy.

While we provide such valuable meta-level insights into what cognitive processes might be most dysfunctional in schizophrenia, our computational modeling approach reached somewhat lower performance in disease classification than certain previous machine-learning studies that were solely focused on prediction performance alone (e.g., Silva et al., 2014). This observation was entirely expected. In statistical data analysis in general, there is a widely recognized tension between predictive performance and model interpretability (Bishop, 2006; Bzdok and Yeo, 2017; Hastie et al., 2009). Machine-learning algorithms are particularly suited to achieve highly accurate predictions in a brute-force fashion, which is why they might be promising for precision psychiatry (Bzdok and Meyer-Lindenberg, 2018; Chekroud et al., 2017). However, such purely data-driven approaches were sometimes criticized for offering less direct insight into the cognitive or neurobiological architecture of schizophrenia. Acknowledging the often-incompatible goals of revealing the underpinnings of a disease and best-possible bare prediction outcomes, our study prioritized the interpretability of the statistical framework. The introduction of condensed neuroscientific knowledge in a principled fashion enabled the study of compromised cognitive processes in a major psychiatric disease. The compression of the original brain information into interpretable summaries came at the expense of best-possible prediction accuracy (Hastie et al., 2009; Kuhn and Johnson, 2013; Shmueli, 2010). Additionally, the classification performance of our predictive model might be compromised by using a multi-site dataset. Such samples often introduce additional sources of variance that differ across sites (e.g., scanner software) (Dansereau et al., 2017; Nielsen et al., 2013). Nevertheless, clinical samples with patients from several sites are often more representative of the general population and are hence more likely to provide clinically relevant and reproducible results (Abraham et al., 2017; Costafreda et al., 2008).

A further strength of our computational neurocognitive assay lies in leveraging accumulated neurobiological knowledge of executed behaviors and their recruited neural systems to build a single-patient prediction framework (Arbabshirani et al., 2017; Huys et al., 2016; Stephan et al., 2017). Such integration of data-driven and theory-assisted analysis tactics allowed us to discover the most impaired cognitive domains in a rich schizophrenia sample while relying on minimal additional statistical, pathophysiological, and neurobiological assumptions. In this way, our study exemplified *hypothesis mining* on which components of human cognition might be particularly affected in a particular psychiatric disorder. These most promising candidate neural systems can provide a well-founded basis for explicit hypotheses testing on multiple levels of observations, such as genomics, neurotransmitters, and neuropharmacology. We are optimistic that such computational psychiatry investigations could be readily extended to many brain disorders and potentially inform priority agendas in health research (Oquendo et al., 2012).

## Methods

### Cognitive description system: BrainMap taxonomy

The cognitive science community has not yet agreed on a consensus definition for mental categories that broadly make up human thinking (cf. Fox et al., 2005; Poldrack and Yarkoni, 2016). Among other possibilities, the BrainMap initiative provides an established means to describe the repertoire of mental operations (Fox and Lancaster, 1994). Experts have steadily refined the description system over two decades, and today it is one of the most frequently applied taxonomies in research practice (Fox and Lancaster, 2002; Laird et al., 2009a). In particular, BrainMap offers two distinct taxonomies to partition human mental activity into compartments: (1) mental domains that span sensory, motor, affective, and higher-level cognitive processes that are recruited during psychological paradigms (i.e., ‘behavioral domains’) and (2) the types of experimental task paradigms used to evoke cognitive processes of interest in a controlled fashion (i.e., ‘paradigm classes’) (Laird et al., 2009b). Both taxonomic catalogues have been used to systematically annotate >16,000 archived neuroimaging experiments from peer-reviewed publications (Fox and Lancaster, 2002; Laird et al., 2011). The completeness and correctness of the labeling of the neuroimaging experiments has been verified by several members of the BrainMap team. The present study capitalized on both description systems to increase the chances of identifying the most pertinent brain-behavior mappings in schizophrenia. To avoid conceptual overlap between the psychological categories considered within each taxonomy, we removed the hierarchical dependence between mental domains by excluding any top-level classes. For example, we excluded ‘emotion’ as an overarching category, and instead considered the subordinates ‘disgust’, ‘fear’, ‘happiness’, and ‘sadness’. We also disregarded rarely used cognitive concepts, defined as those with less than 50 functional neuroimaging experiments in the BrainMap database. By considering only cognitive domains that can be based on a sufficient number of neuroimaging experiments (Bossier et al., 2018; Eickhoff et al., 2016), we could construct robust and meaningful brain-behavior maps, as we will describe in detail in the next section. A final set of 34 mental domains (Fig. 2) and 50 experimental tasks (Fig. 2; Supplementary Fig. 1) from BrainMap was submitted to computational machine-learning assays to test for their utility in schizophrenia research.

### Constructing cognitive meta-priors in healthy participants: ALE meta-analysis of BrainMap taxonomy

To link brain and behavior in typical neurobiology, we carried out quantitative meta-analysis for the synthesis of existing neurobiological knowledge across tens of thousands of neuroimaging experiments from healthy individuals. For each particular cognitive domain, we derived one whole-brain signature of neural activity changes by using state-of-the-art coordinate-based meta-analysis. The widely-used activation likelihood estimation (ALE) approach established convergence of the peak activations reported by functional imaging experiments (Eickhoff et al., 2012; Eickhoff et al., 2009; Turkeltaub et al., 2012). ALE meta-analysis treated the reported coordinates of significant experimental neural response as centers of 3D probability distributions that capture the spatial uncertainty of neuroimaging results (Eickhoff et al., 2012; Eickhoff et al., 2009; Turkeltaub et al., 2012). The spatial extent of the Gaussian probability distribution incorporated empirical estimates of between-template and between-participant variance of neuroimaging peaks (Eickhoff et al., 2009). For each BrainMap experiment, the probability distributions of the reported peak coordinates were merged into a modeled activation map. The use of a non-additive approach prevented local summation effects (Turkeltaub et al., 2012). Finally, all activation maps associated with a particular cognitive process were united to a probability map. The resulting ALE scores yielded the probability of increased neural activity measured during a particular experimental study for each grey-matter voxel. Since ALE scores are influenced by the number of experiments that they are based on, the meta-analytic networks were z-scored by mean-centering to zero and unit-variance scaling to one. This normalization step of each meta-analytic network aimed at improving the comparability between different cognitive meta-priors. Thus, in both BrainMap taxonomies, meta-analytic literature synthesis provided statistically defensible neural activity maps that quantify the convergence zones for each cognitive category.

### Clinical brain-imaging resources: multi-site schizophrenia cohort

Given the well-documented diversity of schizophrenia symptoms, we evaluated the cognitive meta-priors in a high number of patients from several psychiatric hospitals. All predictive modeling analyses were conducted in a multi-modal imaging dataset combined from different brain-imaging laboratories in Europe and the USA that acquired brain scans from patients with schizophrenia and matched healthy controls (n=428). Written informed consent for study participation was obtained from all participants. The data acquisition was approved by the ethics committees of the universities of Aachen, Albuquerque, Göttingen, Utrecht, and Lille. All patients were diagnosed by board-certified psychiatrists according to ICD-10 or DSM-IV-TR criteria. In healthy controls, any history of neurological or psychiatric disorders was ruled out via structured clinical interview. The dataset included (i) demographic indicators including age and sex, (ii) structural brain-imaging data (sMRI), and (iii) resting-state functional brain-imaging data (fMRI). All behavioral and brain-imaging information was anonymized. The sMRI and fMRI data were acquired on common 3T scanners (see Supplementary Table 1 for details). Preprocessing of the imaging data was performed in SPM8 (Statistical Parametric Mapping, Wellcome Department of Imaging Neuroscience, London, UK, http://www.fil.ion.ucl.ac.uk/spm/) using MATLAB R2014a (Mathworks, Natick, MA). In our analyses, we included only those participants for whom both sMRI and fMRI data were available to be able to jointly examine neurobiological impairment in brain structure and brain function. The final sample of 161 patients and 163 healthy controls presented the basis for our machine-learning workflow (see Table 1 for sample characteristics). The total 324 participants were matched for age and sex, both within and across sites (Table 1).

### Brain structure: voxel-based morphometry

To investigate how brain anatomy in healthy controls deviates from brain anatomy in patients with schizophrenia, we relied on volume information in T1-weighted brain scans (Supplementary Table 1). The preprocessing of the whole-brain morphometric maps was performed using standard settings in the VBM8 toolbox (https://dbm.neuro.uni-jena.de/vbm). The anatomical maps were spatially normalized to MNI space (ICBM-152 template) using the DARTEL toolbox including both affine and non-linear spatial transformation. We then quantified the probability of each voxel to belong to gray matter, white matter, and cerebrospinal fluid to segment the volumetric brain maps into the three tissue types. To remedy bias-field inhomogeneities, we applied a unified segmentation (Ashburner and Friston, 2005). Partial volume correction was carried out to account for blurring into neighboring voxels. Furthermore, nonlinear modulation adjusted for inter-individual volumetric differences during the warping process to MNI space. In this way, we obtained gray-matter volume measures for each participant that were corrected for individual brain size. Additionally, we accounted for potential confounding effects of age, sex, and sites to discourage the predictive algorithms from picking up discriminative information in the brain volumes from these influences of no interest.

### Brain function: intrinsic resting-state connectivity

We examined group differences in neural activity fluctuations in the absence of a controlled experimental task based on fMRI maps acquired by resting-state echo-planar imaging (Supplementary Table 1). Before recording of the task-independent BOLD signal scans, the participants were asked to lie still in the scanner and let their minds wander without thinking of anything in particular. Adherence to the instructions was verified in post-scan interviews. The first four MRI scans of each participant were discarded to account for magnetic field saturation. The resting-state time series were aligned by a two-pass procedure to account for participant movements during the scanning session. After co-registration, the functional resting-state maps were spatially registered to MNI standard space (ICBM-152 template), analogous to the sMRI scans. The resulting brain maps were smoothed by a 12-mm FWHM Gaussian kernel. To further account for potential confounding effects through head motion, we corrected the time series of each voxel by a common set of 24 motion parameters: (i) the six motion parameters extracted from image realignment, (ii) their first derivatives, and (iii) the respective squared terms of original motion parameters and derivatives. This specific motion correction procedure was found to improve ensuing functional connectivity analyses by yielding more specific and sensitive brain signals (Chai et al., 2012; Satterthwaite et al., 2013). We did not apply global signal regression (Murphy et al., 2009; Yeh et al., 2015). The BOLD time series were band-pass filtered for frequencies between 0.01 and 0.08 Hz using the frequency-domain filter in the CONN toolbox (https://www.nitrc.org/projects/conn). This frequency range is commonly assumed to represent neural activity and to be less prone to physiological artifacts such as respiration and heart rate (Fox and Raichle, 2007; Lu et al., 2007). Additionally, the BOLD signal time series of each voxel were converted to z-scores in each participant to allow for group analyses. At the across-participant level, we finally helped remove potential confounding influences by accounting for age, sex, and site differences in the fMRI data.

### Sampling neurobiological characteristics of the brain: complementary data-extraction pipelines

The derived cognitive meta-priors guided information extraction from structural and functional brain measurements by focusing on different neurobiological characteristics in the brain data. We wished to relax *a priori* assumptions on the most relevant principle of brain organization in schizophrenia (Weinberger and Radulescu, 2016). We therefore applied different sampling procedures that reflect common approaches to reduce high-resolution MRI measurements to their essence. It is important to appreciate that these preprocessing steps were critical to ensure that the various cognitive meta-priors yielded the same number of variables in each pipeline to ensure statistical comparability in schizophrenia classification (Hastie et al., 2001). Otherwise, different model complexities of the deployed learning algorithms could have made it difficult to attribute lack of predictability to either the brain data themselves or possible discrepancies of the modeling procedure.

Three overarching strategies profited from distinct and complementary ways to aggregate neurobiological information:

1. *Mining peak locations:* Our “peak activation” approaches concentrated on the most important voxel groups of a given cognitive meta-prior. Target voxels were extracted by searching for locations with the highest probability of increased neural activity during the engagement of a particular cognitive process. Hence, this simple strategy selected a subset of the most important grey-matter voxels from the brain maps guided by the meta-priors. Because the procedure was based on inspection of single voxels in structural or functional brain data, the analyses were perhaps closest, in character, to mass-univariate analyses prevalent in neuroimaging:

a. The “highest absolute activation peaks” approach sampled the voxels with the highest probability of increased neural activity from each cognitive meta-prior without imposing additional assumptions. The original meta-analytic maps of the corresponding neurocognitive priors were used to extract the voxels of generally largest signal changes.
b. The “highest specific activation peaks” approach sampled the peak voxels after accounting for voxels with high across-domain baseline activity after subtracting the mean neural activity level for each voxel across meta-priors. For instance, large-scale analyses showed regions of the saliency network and of the fronto-parietal network to have the highest task-response probabilities (Nelson et al., 2010; Yarkoni et al., 2011). The preference of voxels that were specifically increased in neural activity by a particular cognitive process tended to enhance the relative differences between meta-priors.
c. The “standardized activation peaks” approach accounted for both the mean and the variance of each voxel observed across meta-priors. Before identifying the target voxels, we subtracted the mean neural activity and scaled the voxels to unit variance across meta-priors.
2. *Mining regional characteristics:* An alternative procedure was deployed to derive new biologically meaningful region variables to acknowledge the perhaps still most dominant view on brain organization (Kanwisher, 2010; Passingham et al., 2002). Our “regional specialization” approaches were faithful to the idea that the brain is partitioned into a mosaic of localized, non-overlapping territories (cf. Finn et al., 2015; Glasser et al., 2015). The perspective emphasizes that cognitive processes may be realized by recruitment of neuronal populations that occur in disjoint brain compartments. Clustering methods naturally dovetail with grouping similar voxels into distinct brain regions (Eickhoff et al., 2015; Thirion et al., 2014) by assigning each voxel to exactly one brain region only. Three complementary clustering algorithms were used to merge voxels to homogeneous clusters such that the voxels within a region are more similar to each other in structural or functional properties than to voxels from other regional territories:

a. K-means clustering iteratively readjusts the territory centers and then re-assigns the voxels to each nearest cluster hotspot by minimizing the Euclidean distance of the voxels within each cluster (Lloyd, 1957; Nanetti et al., 2009). The partitioning procedure relied on minimal assumptions and imposed, for instance, no spatial constraints so that the extracted regions were not necessarily spatially contiguous.
b. Ward clustering is a hierarchical clustering algorithm that successively combines the most similar voxels until a number of specified regions is reached. Ward clustering aims at minimizing the variance between voxels within each cluster (Johnson, 1967). In contrast to the more liberal constraints of k-means, only neighboring voxels were fused which resulted in spatially contiguous regions in the brain (Abraham et al., 2014).
c. Spectral clustering transforms the data in a non-linear fashion, which complements the k-means and ward clustering approaches. The non-linear transformation enabled the spectral clustering algorithms to discover non-convex clusters that contrasted with those obtained with k-means and ward clustering. First, a similarity graph was constructed that represented spatial proximity between the voxels (van Luxburg, 2007). Then, the graph was partitioned such that the weight of the edges cut was small compared to the weights of the edges inside each cluster (Donath and Hofman, 1973; Thirion et al., 2014). Different from the k-means clustering approach and analogous to the ward clustering approach, only spatially contiguous voxels were merged into region clusters.
3. *Mining network characteristics:* Yet another complementary procedure accommodated the organizational perspective of brain function arising from an interplay of distributed, overlapping networks (cf. Smith et al., 2009). Our “distributed networks” approaches created network variables by focusing on the functional connections between distinct brain compartments that are cross-regionally integrated (Sporns, 2014; Van Essen et al., 1992). This conceptualization is naturally captured by matrix decomposition algorithms that broke down the brain into a number of hidden distributed network components (Smith et al., 2009). In contrast to the “regional specialization” approach, each voxel belonged to each of the components to varying degrees:

a. Principal component analysis (PCA) is a widespread procedure that searches for spatially uncorrelated network components that explain the observed variance distributed in the brain data (Shlens, 2014). The orthogonal components consisted of linear combinations of the voxels, while all grey-matter voxels were assigned continuously to each network and nonlinear relationships between the variables were ignored.
b. Sparse PCA is a recent variant of PCA that additionally exploits the fact that often only a subset of voxels is relevant for extracting coherent network components to explain most of the observed variance in the data (Zou et al., 2006). A sparse representation was accomplished by additionally imposing a parsimony constraint (L1 penalty) that also partly relaxed the orthogonality assumption of classical PCA (Chennubhotla and Jepson, 2001).
c. Independent component analysis (ICA) is able to discover the sources of variation that independently contributed to the observations in the brain, instead of imposing uncorrelatedness between networks such as in the PCA approaches (Calhoun et al., 2001; Hyvarinen, 1999). Complementing PCA, the neural signal was non-linearly separated into network components where a particular network node could readily contribute to more than one network component (Hyvarinen, 1999).

In sum, our data preparation pipelines sampled complementary aspects of the brain by considering each participant’s high-dimensional brain scans by means of importantly different dimensionality-reduction techniques. This meta-prior-guided extraction of sMRI and fMRI data enabled direct comparison of our analytical approach in (i) brain structure, (ii) brain function, and (iii) their combination. For the joint analyses of both imaging modalities, we concatenated the extracted sMRI and fMRI data for each participant. In sum, nine complementary sampling pipelines extracted meaningful neurobiological information from patients and controls by guidance through the meta-priors and commonly used dimensionality-reduction techniques.

### Confederating ensembles of cognitive meta-priors: model stacking for integrated prediction

For each cognitive meta-prior, we used the extracted brain information for training a predictive pattern-learning algorithm (i.e., ‘base model’) to distinguish healthy controls from patients with schizophrenia. The collection of ensuing predictive models was subsequently incorporated into one higher-level predictive model (i.e., ‘composite model’) to stratify the cognitive domains according to their forecasting performance. This two-step stacking strategy (Breiman, 1996; Wolpert, 1992) was a native choice to put each meta-prior to a comparable scale and identify their relative relevance for schizophrenia classification, despite their naturally diverging neurobiological representations. By placing all meta-prior models on a common scale for each of both taxonomies, the summary model automatically ranked the whole set of cognitive processes according to their potential involvement in schizophrenia.

1. Base models: Separately for each cognitive category of a particular taxonomy, we fitted one simple linear classification model to disambiguate the groups based on the extracted and z-scored brain data. In analyses involving fMRI, the 25^th^ percentile of highest scoring resting-state connectivity features was selected first (in the training data, cf. below) according to the strength of univariate relationships with the participant group. The adaption of this feature-selection step, similar to the sMRI analyses, was intended to improve comparability between both imaging modalities. For each cognitive meta-prior, we thus fitted a logistic-regression algorithm to the extracted sMRI and fMRI data of a larger part of the participants (i.e., training sample). Then, we used the logistic regression to predict disease state (schizophrenia versus health) in the previously left-out participants (i.e., test sample). The evaluation of disease state in new participants yielded practically relevant predictions because the algorithm did not visit the participants during model estimation (Gabrieli et al., 2015). Thus, the base models predicted the probability to be affected by schizophrenia from the structural and functional brain data. The independent disease state predictions of each cognitive meta-prior served as input for the integrated model.
2. Composite model: The meta-prior specific predictions of the base models were combined for training a more elaborate predictive model that considered the separate relevances of all cognitive processes of a taxonomy at the same time for schizophrenia detection. For this purpose, we used a random-forest algorithm because the high-capacity classifier is susceptible to complicated nonlinear relationships combined with the possibility of model interpretability. This pattern-learning algorithm involves fitting a collection of decorrelated decision trees and uses their majority vote for prediction (Breiman, 2001; Louppe, 2014). As a first advantage, the ensuing committee classifier was able to quantify the single meta-priors regarding their contribution for schizophrenia classification (Breiman, 2001; Louppe et al., 2013). As an ensuing second advantage, random forests could uncover potential non-linear interactions between the meta-priors, and thus their corresponding cognitive classes. Since we wished to reduce variability in the classification process, we set a common choice of trees in the random forest to 1000. The depth of the trees was set to 5 because higher-order interactions between cognitive processes would have evaded ready visualization or interpretation. The maximum number of features considered at each split in a tree was set to 1, which encouraged decorrelated trees and further improved the equal opportunity between the meta-priors. Overall, the stacking of meta-prior-specific base models into a summary model enabled us to rank each taxonomy of cognitive domains in their ability to distinguish between patients and controls.

### Model evaluation: nested 10-fold cross-validation for single-participant prediction

For the obtained domain-overarching predictive models we quantified the capability to correctly distinguish brain data from participants that we would observe in the future. To examine the performance of the neurobiologically informed composite model in participants whom the algorithm has not seen before, we implemented a nested, stratified 10-fold cross-validation. The participants were divided into ten balanced data splits (folds), each preserving the percentage of participants of both classes. The predictive model was repeatedly fitted on 90% of the data and subsequently assessed in the brain-data of the left-out 10% of the participants (Hastie et al., 2009; Stone, 1974; Stone, 1978). After ten iterations of model fitting and testing, the percentage of correctly classified test participants was averaged across folds. The nested variant of the cross-validation scheme ensured that only actual base model predictions were fed into the composite model as one important characteristic of stacking procedures (Hastie et al., 2009; Wolpert, 1992). Note that we aimed at validating the practical plausibility of our approach for single-patient prediction, instead of tuning our model towards high prediction accuracies. The obtained quantity yielded the crossvalidated prediction accuracy of the integrative composite model to generalize to future participant samples from the population.

To additionally evaluate how much the obtained classification performances in other schizophrenic patients would be expected to vary, we computed their 95% population confidence intervals using bootstrapping (Efron and Tibshirani, 1994). The statistical procedure generates alternative datasets we could have obtained from the sample that we had by repeatedly drawing random samples of the original data with replacement. In 1000 bootstrap iterations, the identical nested cross-validation scheme was carried out on the perturbed participant samples. This uncertainty interval estimation answered how the classification success was expected to vary in the broader schizophrenia population. The classification performance of the composite model enabled comparisons between (i) different imaging modalities (sMRI, fMRI and combined sMRI and fMRI data), (ii) the set of complementary procedures of neurobiological sampling, and (iii) two BrainMap taxonomies (mental domains and experimental tasks). Furthermore, the model performance allowed us to validate the composite model against models not informed by cognitive meta-priors (cf. below). In sum, testing the generalizability of the composite model helped us to gain confidence in the robustness and potential clinical usefulness of our data-analysis approach.

### Model inspection: variable importance, nonlinear effects, and predictive relevance maps

We explored the predictive contribution of the individual meta-priors in a particular taxonomy in an identical process. Our stacking-model approach allowed reverse engineering which 34 mental domains or 50 experimental tasks might be most affected in schizophrenia. For this purposes of model interpretation, the composite model was initially refitted on the full participant sample (Hastie et al., 2009). Random forest algorithms naturally afford a quantitative measure of relative importance for each input variable (Breiman, 2001). Technically, the variable importance of a meta-prior provide a convenient summary of the mean decrease in the misclassification rate across all branch splits in which a specific variable was used in a grown decision tree to separate the healthy and schizophrenic group (Louppe et al., 2013). Since each input variable fed into the random forest corresponded to a single meta-prior, the variable importance of the composite model weighted the ensemble of cognitive domains in a same step. We capitalized on these relative importance weights to assign each meta-prior a ranking position according to their disease discriminability. The highest values of importance indicated the first rank. To quantitatively estimate the precision of the ranking positions of the meta-priors in the general population, we estimated the 95% confidence intervals in 1000 bootstrap iterations by repeatedly fitting the final model to resampled alternative datasets that we could have been freshly drawn from the population. Since the model evaluation revealed that none of the nine pipelines were uniformly superior, we averaged the ranking positions across all of them to enhance impartiality of neurobiological assumptions. As an overall uncertainty estimate accounting for random sampling effects, the 95% confidence intervals were calculated from the bootstrapped distributions of the variable importance across pipelines. In addition to the relative brain-behavior contribution in schizophrenia, the random forest also allowed the investigation of potential non-linear interactions between the cognitive meta-priors. As two-way interactions are easier to understand by humans than higher-order interactions, we detailed the interaction surface for each pair of cognitive domains after accommodating the remaining cognitive meta-priors contribution to the prediction of schizophrenia. To additionally capture discriminative characteristics of schizophrenia on the neurobiological level, we investigated which brain regions were most pertinent for disease classification. For this purpose, we multiplied each cognitive meta-prior with its respective importance weight of the composite model. Then, we averaged the ensuing domain-maps to provide a global predictive relevance map for schizophrenia. Across neurobiological sampling pipelines, we thus inspected the predictive value of the cognitive domains, their interplay to algorithmic modeling of the group of each participant, and their neurobiological basis.

### Testing the cognitive specificity of schizophrenia predictability: comparison to a null model

A negative test ensured the *fit for purpose* of the final predictive model across imaging modalities (Kuhn and Johnson, 2013). This sanity check answered the question ‘Did we successfully distinguish patients from controls because the summary model captured the individual configurations of cognitive facets rather than other characteristics of our participant sample?’. To this end, we examined the null hypothesis that no coherent relation exists between the spectrum of cognitive involvements of healthy controls and schizophrenia patients. The placebo hypothesis was put to the test by a non-parametric permutation procedure (Efron, 2012; Winkler et al., 2016). We specifically corrupted cognition-related structure in the data, while leaving the other joint probabilities intact. That is, we only perturbed variance in the data related to the alternative hypothesis of individual expressions of cognitive meta-priors achieving disease classification. We randomly exchanged the importances of individual meta-priors between participants, separately in patients and controls, before they were fed into the composite model. This permutation scheme preserved the manner in which each cognitive meta-prior scored and the disease structure of our sample. Yet, the procedure was targeted at altering the participant-level pattern of meta-prior expressions. Put differently, the permutation changed how the meta-prior relevance co-occurred in combinations within patients and within controls. Based on these slightly permuted data, the same composite model was fit 1000 times to compute a distribution of classification performances that occur under the null hypothesis. Subsequently, we compared the actually obtained prediction accuracy of our meta-model against the no-effect distribution. In each data analysis pipeline, the comparison of the composite model to the performance of a null model allowed us to ascertain that our disease classification was based on the combined cognitive facets in individual participants.

### Scientific-computing implementation

Our data-processing workflow was implemented in Python 2.7. We chose the open-source programming language to enable the reproducibility of our results and encourage reuse of our code in future projects. All computational analyses relied on unit-tested implementations of state-of-the art machine-learning algorithms as provided by scikit-learn 18.1 (Pedregosa et al., 2011). The application of the predictive models to high-dimensional neuroimaging data was facilitated by nilearn 3.0 (Abraham et al., 2014). The full analysis workflow completed after >7 days on our computing cluster hosted at the Rechenzentrum of RWTH Aachen University with 52 cores and 512 GB working memory. All data analysis scripts are publicly available for transparency and reuse at: https://github.com/TMKarrer/domain-ranking-scz.

## Acknowledgements

Teresa M. Karrer is funded by a full PhD scholarship of the German National Merit Foundation. Dr. Bzdok is funded by the Deutsche Forschungsgemeinschaft (DFG, BZ2/2-1, BZ2/3-1, and BZ2/4-1; International Research Training Group IRTG2150), Amazon AWS Research Grant (2016 and 2017), as well as the START-Program of the Faculty of Medicine (126/16) and Exploratory Research Space (OPSF449), RWTH Aachen. Dr. Bassett acknowledges support from the ISI Foundation, the Alfred P. Sloan Foundation, the John D. and Catherine T. MacArthur Foundation, and the Paul Allen Foundation.

## Author contributions

D.B. conceived the project. T.M.K. and D.B. conducted the data analysis. T.M.K. completed the first manuscript draft. O.G., G.V., and B.T contributed analytical solutions. B.D., O.G., A.A., and R.J., contributed neuroimaging data. All authors provided feedback and revised the manuscript.

